# An atlas of the Norway spruce needle seasonal transcriptome

**DOI:** 10.1101/2021.07.12.452085

**Authors:** Pushan Bag, Jenna Lihavainen, Nicolas Delhomme, Thomas Riquelme, Kathryn M Robinson, Stefan Jansson

## Abstract

Boreal conifers possess a tremendous ability to survive and remain evergreen during harsh winter conditions and resume growth during summer. This is enabled by coordinated regulation of major cellular functions at the level of gene expression, metabolism, and physiology. Here we present a comprehensive characterization of the annual changes in the global transcriptome of Norway spruce needles as a resource to understand needle development and acclimation processes throughout the year. In young, growing needles (May 15 – June 30), cell walls, organelles etc. were formed, and this developmental program heavily influenced the transcriptome, explained by over represented Gene Ontology (GO) categories. Later changes in gene expression were smaller but four phases were recognized: summer (July-August), autumn (September-October), winter (November-February) and spring (March-April), where over represented GO categories demonstrated how the needles acclimated to the various seasons. Changes in the seasonal global transcriptome profile were accompanied by differential expression of members of the major transcription factor families. We present a tentative model of how cellular activities are regulated over the year in needles of Norway spruce, which demonstrates the value of mining this dataset, accessible in ConGenIE together with advanced visualization tools.

**Significance statement:** The development of Norway spruce needles and their annual cycle of biochemical activities (photosynthesis in the summer, adaptation and survival of the harsh boreal winter) is not well understood. We use deep RNA sequencing to profile the transcriptome over the season, and show how the dataset could be used to give information about “what needles do” over the year, and are here making the dataset available for the scientific community.

**Graphical abstract:** 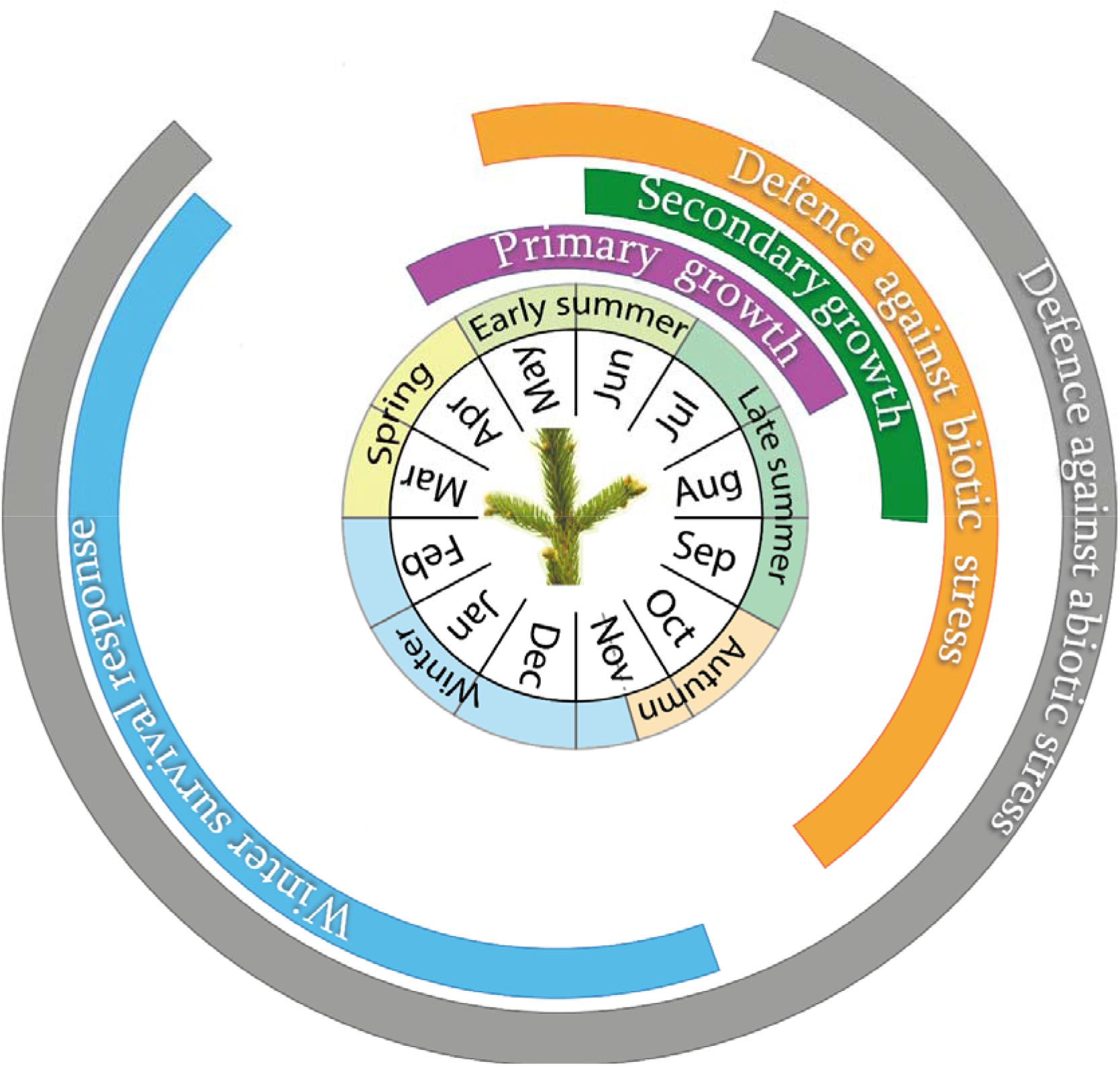

## Introduction

Vascular plants have colonized most land on earth except where the lack of accessible water severely restricts growth, either due to low precipitation or in polar areas where it is not accessible since frozen. In regions close to these habitats, natural selection has caused plants to develop strategies to cope with unfavourable conditions, often by avoidance (*e.g.* annuals that survive as seeds) or by developing strategies that enable them to persist even under very harsh environmental conditions. Conifers (*Pinophyta*) constitute a division within the gymnosperms that branched off from the lineage leading to angiosperms (flowering plants) more than 200 million years ago (Savard et al. 1994, Friedman and Cook 2000). Among the ca. 630 known conifer species in 70 genera (Nystedt et al. 2013), some have evolved remarkable adaptations, most notably the ability to grow under very harsh and dry conditions and to retain their leaves (needles) over several years and particularly during the boreal winter.

Boreal forests in northern Europe are dominated by the evergreen conifers Scots pine (*Pinus sylvestris*) and Norway spruce (*Picea abies*). In this environment, the lifespan of a Norway spruce (*Picea abies*) needle is typically five years or more. It is believed that boreal conifers generally enter a lockdown mode during the winter with very limited biochemical activity (Moreno, Carlson, and Alatalo 1988; Öquist and Huner 2003; Dhuli, Rohloff, and Strimbeck 2014). However, even in the middle of the winter, some activities may be ongoing that allow further acclimation and promote survival strategies during cold periods. Several studies on conifers in the past decades have shed light on this adaptation mechanism. One hypothesis is based on the negative correlation of leaf longevity with leaf nitrogen content (Reich et al. 1995; Diemer 1998; Merry et al. 2017; Jokipii◻Lukkari et al. 2017) but adaptation of photosynthesis (Öquist and Huner 2003) is, of course, also necessary to enable conifers to keep their needles intact during cold periods. In any case, acclimation needs to be triggered well before the harsh conditions appear in order to rearrange cellular metabolism (Harsch et al. 2014) and the annual cycle of activity requires adequate transcriptional regulation to achieve proteomic, lipidomic and metabolomic changes.

Gene expression in conifer needles over the season has been studied previously (Prunier, Verta, and MacKay 2016; Cronn et al. 2017; Jokipii◻Lukkari et al. 2017), but it was not until the first conifer genomes had been sequenced and high-throughput methods such as RNA sequencing had been developed that it became possible to establish comprehensive expression atlases of gene expression. As part of our project to characterize the Norway spruce genome (Nystedt et al. 2013) we sampled needles throughout a year and subjected them to RNA sequencing. Besides utilization in the annotation of the Norway spruce genome, this dataset, representing a wide transcript coverage and highly frequent sampling, was generated to benefit the scientific community, and is made publicly available as a part of the ConGenIE database (https://congenie.org/)(Sundell et al. 2015). Here, we investigate the quality of the dataset and the extent to which conclusions about the annual regulation of gene expression in needles of spruce grown in the field under natural conditions can be drawn from this RNA sequencing analysis. Our study demonstrates, for example, how transcription factors are expressed and altered global transcriptional patterns in the needles regulate adaptation of cellular functions in a coordinated fashion over the seasons, reflecting the acclimation that could account for the survival of Norway spruce needles in harsh boreal environmental conditions.

## Results

### Five phases of gene expression were found over the year

Shoot growth in Norway spruce starts in our boreal climate in May and continues for about a month, and the duration of this growth period and the length of the newly formed shoot is strongly influenced by environmental factors, mainly temperature. In this study, we sampled Norway spruce needles in total 28 times from May 16 one year to May 11 the next year. The weather conditions during the sampling period are summarized in Fig. 1A. The transcriptome data presented here are relative rather than absolute values, to conform with how such datasets typically are visualized; absolute values are available in Table S2.

**Figure 1.**
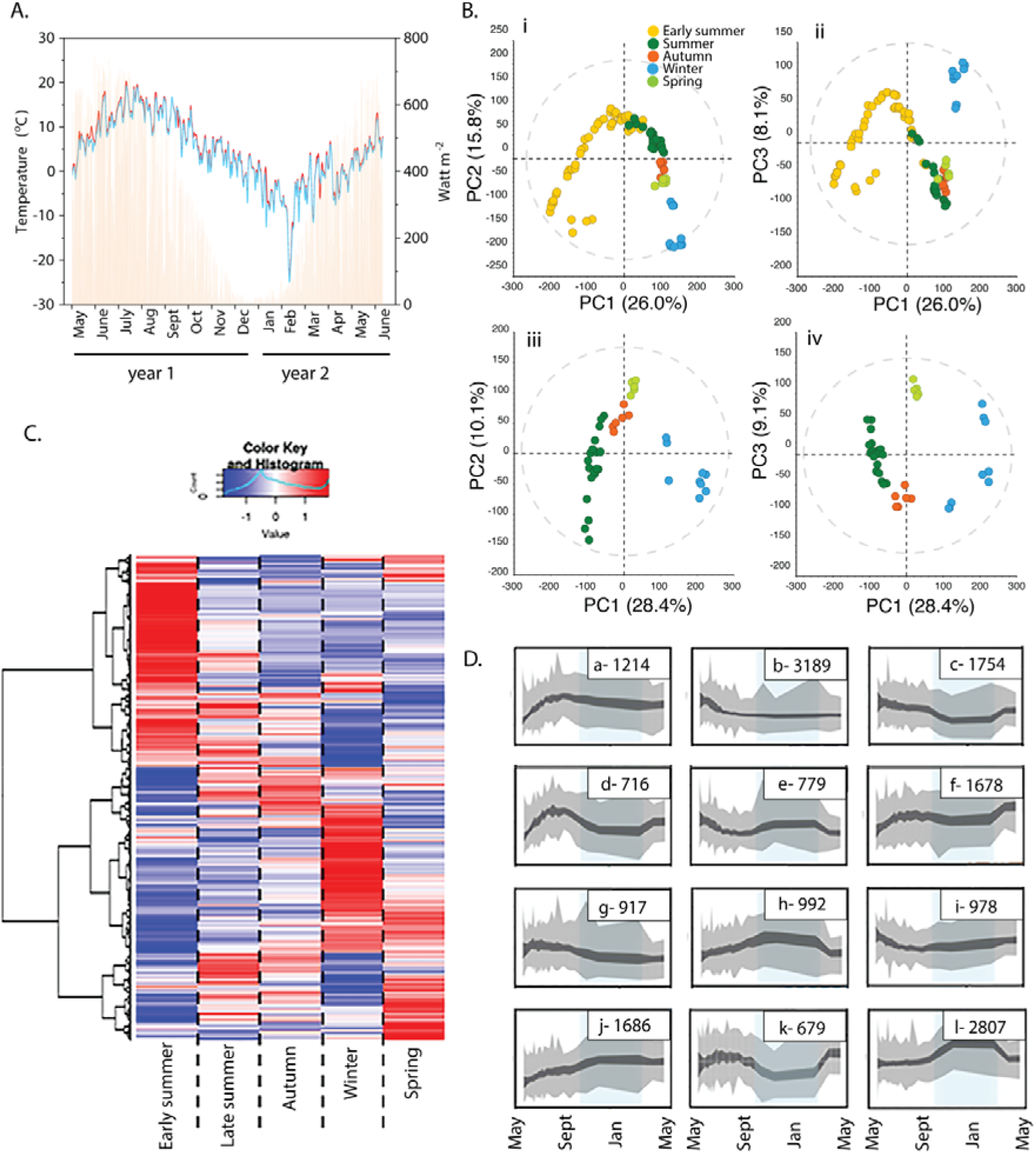
Global transcriptome patterns as affected by seasons and the change in weather parameters over the study period. ◻ A. Daily air temperature (◻ C) (Max in red, and Min in blue) and solar radiation (Watt m^−2^, in orange) recorded on hourly basis over the sampling period from May 2011 to May 2012. B. Principal component analysis (PCA) of transcriptome data during 2011-2012; considering all time points (B-i, B-ii) and excluding early summer (May and June) period (B-iii, B-iv). C. Heatmap of seasonal expression patterns of 28354 nuclear Norway spruce genes, D. The temporal expression profiles of twelve clusters (k=12).

During the period of shoot elongation, needles also develop, so our first samples were very young developing needles. The overall variation in the genome data was examined using Principal Component Analysis (PCA). PCA of the whole dataset yielded eight significant components (95% confidence interval with the cross-validation approach(Eastment and Krzanowski 1982)) where the first component (PC1) explained 26 % of the variation and separated (particularly if combined with PC2 and PC3) samples from May and June from samples collected during the other seasons (Fig. 1B- i, ii). It has been observed in other species, for example in our previous studies on *Populus tremula* (Sjödin et al. 2008), that gene expression during early leaf development is mainly driven by developmental changes as the cellular components/organelles are being synthesized and assembled during this phase. To avoid that these developmental stage-dependent differences in the transcriptome in early summer – where we also had a great proportion of samples – obscured changes that could be coupled to acclimation/other seasonal effects, we first split the dataset into two, samples collected during May and June, hereafter named early summer (ES) and the rest excluding ES. A PCA without ES samples (Fig. 1B- iii, iv) had five significant principal components (95% confidence interval), with the first three components explaining approx. 50% of the total variation. The first two components (PC1, 28.4% and PC2 10.1%, Fig. 1B- iii) separated samples from October-February (hereafter named winter) from July-August (summer) and samples from September (autumn) and April-May (spring) and based on the third component (PC3 9.1%), autumn and spring samples also were separated from each other (Fig. 1D).

An hierarchical clustering analysis (HCA) of global genome profiles – ES samples included – illustrated the distinct expression patterns between the five seasons defined above and 12 major clusters of expression profiles (each with over 500 genes) were identified (Fig. 1D). Four of them (A, B, G and I) varied considerably during early summer but less during the rest of the year, whereas the others showed variation throughout the year. Taken together, both PCA and HCA – including or excluding early summer samples – provided support for a division of the samples into five classes: early summer where gene expression was mainly driven by needle development and summer, autumn, winter, and spring where gene expression related to acclimation was more prominent. As seen in the PCA (Fig. 2), the gene expression patterns shift gradually in early summer, being driven by a developmental process. Although early summer samples were slightly overlapping with summer samples, acclimation periods (summer-spring) stood out separately in the PCA from the samples from May and June. July 1 corresponded fairly well to a transition point in gene expression and to the time when needles have stopped elongating. In the following sections we will use this classification of samples into five seasons to draw conclusions about, firstly, gene expression during needle elongation and development and, secondly, acclimation of the needles to changing seasons. The complete dataset is however available in ConGenIE to facilitate alternative, independent analyses using its visualization tools, or for downloading.

**Figure 2.**
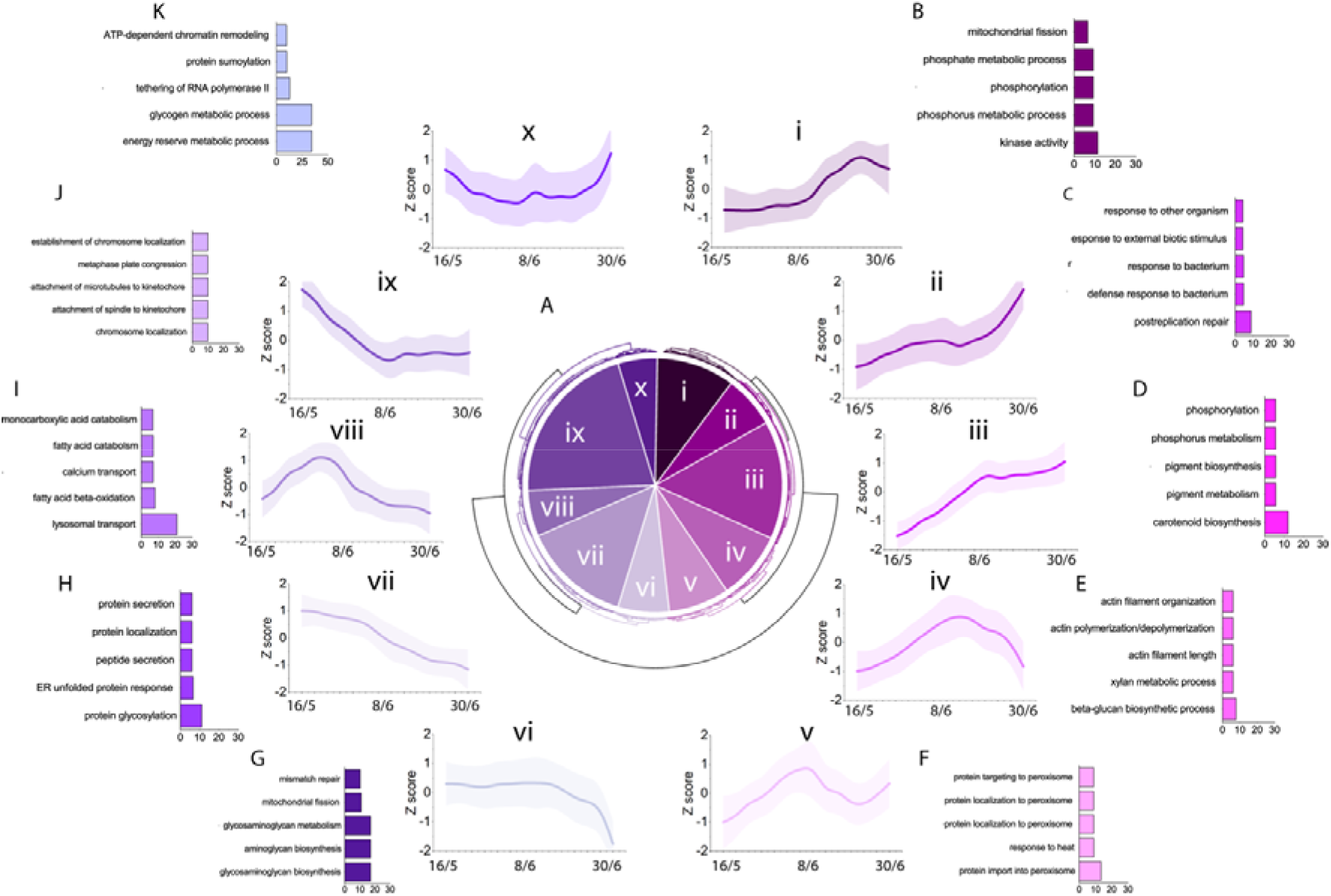
Hierarchical clustering analysis of genes in early summer, cluster patterns and enriched Gene Ontology (GO) terms for biological processes and cellular components. ◻ A. Ten temporal patterns (i to x) obtained from hierarchical clustering of genes (tree cut off h=10). B. to L. Top 5 enriched GO terms for biological processes (BP) in each gene cluster (i to x) are shown. See complete list of GO terms in Suppl. Table 3.

### The transcriptome profile of early summer needles reflected developmental processes

An HCA of the global genome (GT) separated the ES samples alone (GT-HCA) into ten major clusters (Fig. 2) with distinct expression profiles, five of them (Fig. 2 i-v) with overall higher expression levels in June and five (Fig. 2 vi-x) with higher levels in May. We also performed a PCA using only genes (ca 8500) that were first identified as differentially expressed (DE) in ES, compared to the other seasons (DE-HCA), and here six major clusters were identified (Figure S2). Some of the clusters identified in GT-HCA matched clusters identified in DE-HCA (such as cluster ix in the GT-HCA and cluster iii in DE-HCA, Figure S2 L) while, for example, cluster i in GT-HCA contained genes from both clusters i and ii in DE-HCA. To confirm that these clusters corresponded to developmental phases where genes involved in certain biological processes were expressed, we performed a GO term enrichment analysis of clusters from both HCAs (Table S3). Clearly, the most enriched GO terms corresponded to activities expected to occur at certain stages. For example, cluster ix in the GT-HCA, which had highest expression in the earliest samples, was most enriched in GO terms related to cell division (e.g. chromosome localization, attachment of spindle to kinetochore, metaphase plate congression). Cluster iii, most expressed in the earliest samples in the DE-HCA, was enriched for GO terms including cell cycle, DNA replication, prophase and M phase. Clusters with the highest expression in June were enriched with other GO terms, for example cluster ii in GT-HCA in secondary metabolite biosynthesis and oxylipin biosynthesis, cluster i in DE-HCA in programmed cell death (PCD) involved in cell development, monoterpene biosynthesis and lignan biosynthesis and iii in DE-HCA in phosphorus metabolism and pigment biosynthesis. From these results it was clear that this dataset was useful to visualize the developmental program in the newly formed needles, from cell division and through cell expansion to organelle development in May, to secondary cell wall synthesis and activated defence against other organisms in June. As early needle development is outside the scope of this contribution, we do here not further analyse gene expression in ES needles.

### Different biological process GO terms were enriched among the four seasons

To focus on seasonal acclimation, we performed an analysis of differentially expressed genes among the four seasons summer, autumn, winter and spring. Almost 9000 genes showed changes in their expression levels between consecutive seasons (Fig. 3A, Table S4). The biggest differences were found between autumn and winter and winter and spring with different dynamics of up- and downregulation of genes. The autumn to winter transition involved a large number of downregulated genes whereas winter to spring transition was associated with a large number of upregulated genes. An HCA of the expression levels of differentially expressed genes across the seasons displayed seven temporal patterns (Fig. 3B). These could broadly be described as upregulation in spring and downregulation in autumn (cluster I), downregulation in spring and upregulation in autumn (II), upregulation in winter (III), downregulation in winter (IV), upregulation in spring and summer and downregulation in autumn and winter (V), and upregulation specifically in summer with two distinct patterns with either consistent high expression (VI) or gradually decreasing expression through summer (VII). The overlap in differentially expressed genes among seasons (Venn diagrams, Fig. 3C) was limited, and corresponded with the distinct seasonal profiles observed in the PCA (Fig. 1B iii, iv).

**Figure 2.**
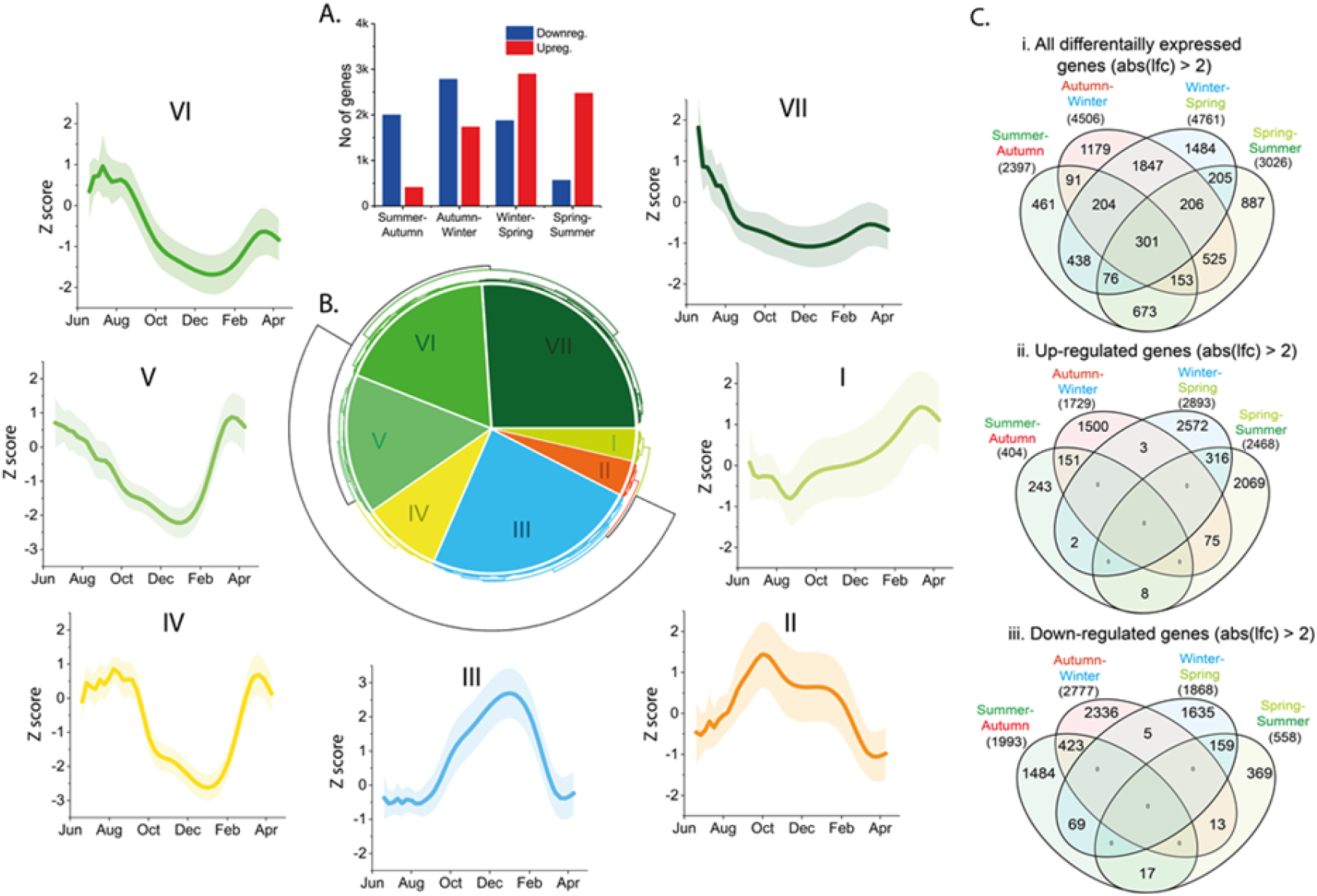
Clustering analysis and the temporal patterns of differentially expressed genes over the four seasons. ◻ A. The number of up- and down-regulated genes between consecutive seasons (DESeq2, log_2_ fold change >2, FDR-adjusted p-value <0.01); B. Seven clusters and their temporal patterns (I-VII) obtained using hierarchical clustering analysis (tree cut off h=10); C. Venn diagram showing the overlap in differentially expressed (C-i), up- (C-ii) and down-regulated (C-iii) genes between consecutive seasons, total number of differentially expressed genes are shown in parentheses. See complete clustering and DE list information in Suppl. Table 4.

Gene Ontology (GO) term enrichment analyses were performed for each cluster (Table S5) and enriched GO terms in the clusters were integrated in a network to give an overview of the connections between altered biological processes through the seasons and to identify potential processes affected by a specific season (Fig. 4). Biological processes predominantly enriched in summer were involved in cell wall processes, lignin biosynthesis and leaf morphogenesis (Fig. 4, cluster VI, VII), reflecting that secondary cell wall synthesis continues after shoot/needle elongation has ceased. Defence response related GO terms were also enriched, presumably to allow needles to withstand abiotic and biotic stresses (Fig. 4, cluster VI, VII).

**Figure 4.**
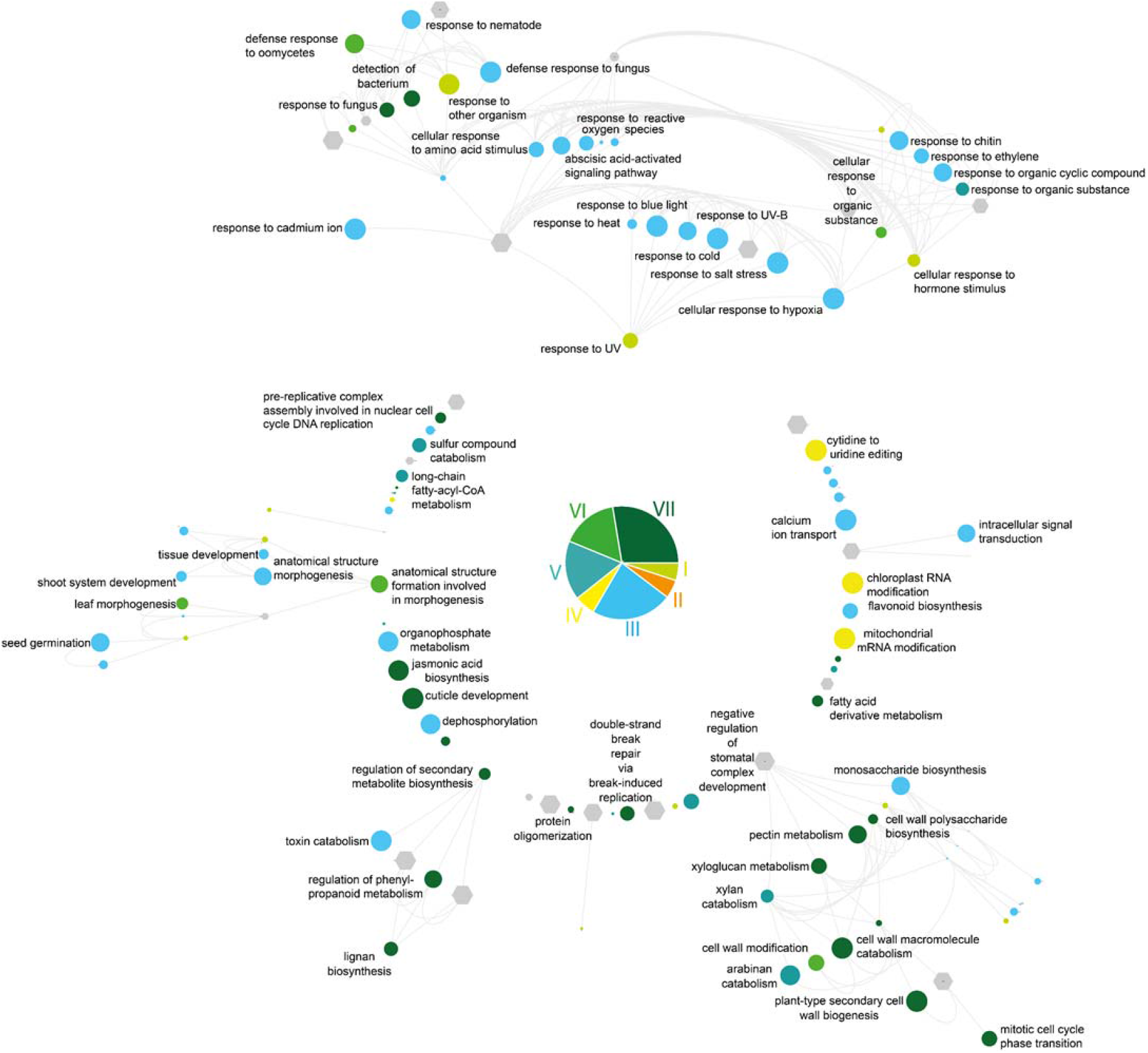
Gene Ontology (GO) term network of enriched biological processes in gene clusters affected by seasons. ◻ Colour of the nodes corresponds to the seven temporal clusters in Figure 3. Grey nodes indicate GO terms that were enriched in multiple clusters. Size of the node is proportional to the (−)log_10_(P) value of enrichment significance; larger node size denoting higher significance. See complete list of GO terms in Suppl. Table 5.

### Winter and autumn transcriptomes indicate upregulation of stress responses and down regulation of cellular activity

Biological processes upregulated in winter included several stress responses such as response to cold, water stress, reactive oxygen species and hypoxia, many of which involve overlapping genetic regulation and phytohormone signals, abscisic acid (ABA) and ethylene (Fig. 4). As we are particularly interested in the overwintering capacity of needles we visualized all significant GO terms associated with clusters III (Fig. 5) and IV (Fig. 6) that displayed enhanced or repressed expression in winter, respectively. Downregulated genes in winter were specifically enriched in chloroplast and mitochondrial RNA processing including genes encoding multiple organellar RNA editing factor (MORF) proteins and pentatricopeptide repeat (PPR) proteins (PPRPs) (Fig. 5). Most of the detected PPRPs belonged to the subfamily of plant combinatorial and modular proteins (PCMPs) that function in cytidine to uridine editing of RNAs (Fig. 5). PPRPs and MORFs form RNA editing complex, where PPRPs are putative to bind plastid RNA whereas MORFs selectively bind to PPRPs.

**Figure 5.**
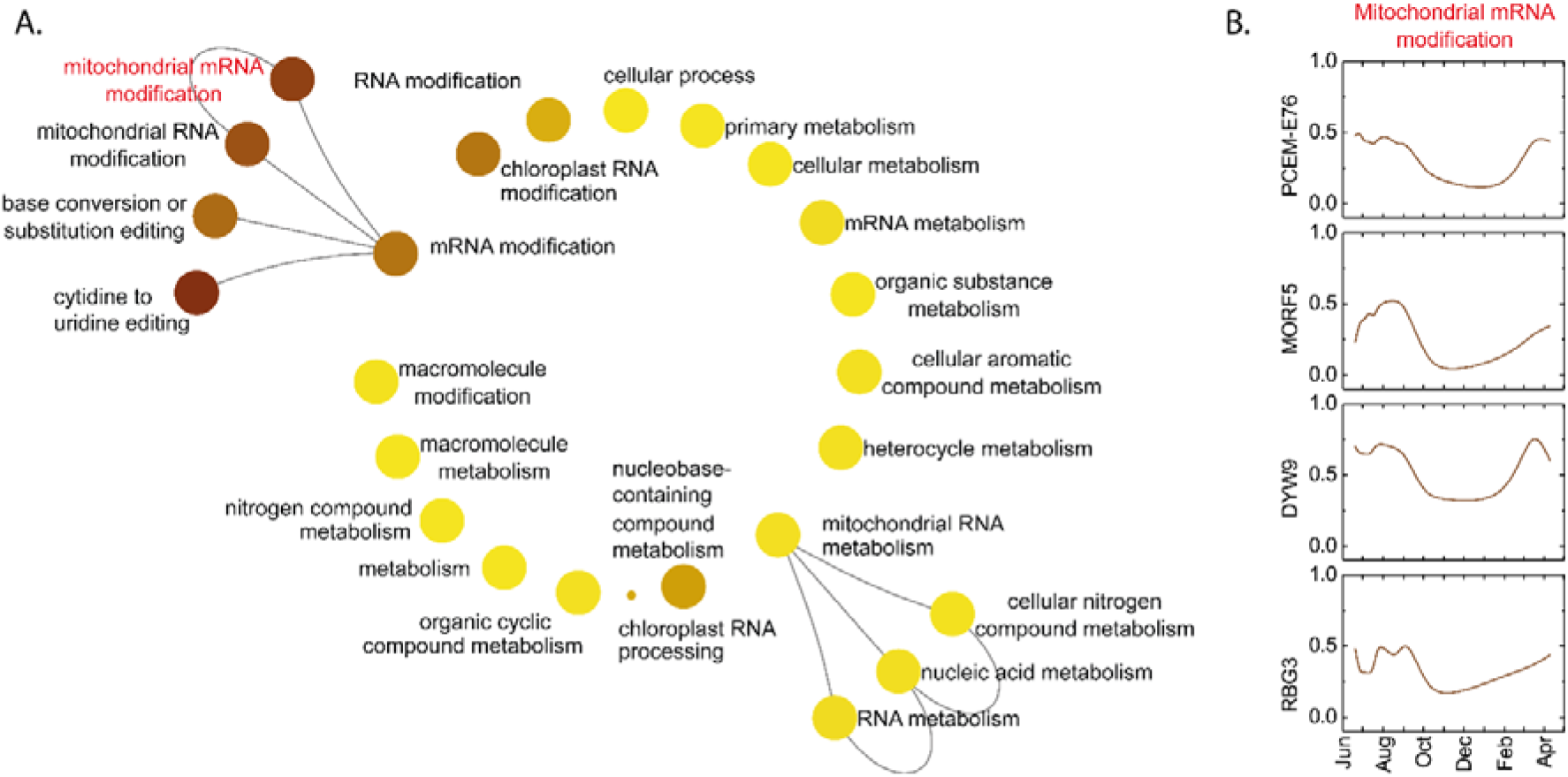
Gene Ontology (GO) term enrichment analysis of the genes in the yellow cluster (IV) downregulated in winter and the expression patterns of selected genes related to the enriched processes. ◻ A. Enriched GO terms for biological processes in yellow cluster (IV). Size of the node is proportional to the (−)log_10_(P) value; larger node size denotes higher significance and darker colour denotes higher fold enrichment of the GO term; B. Temporal expression patterns of PCEM, MORF5, DYW9 and RBG3 related to mitochondrial mRNA modification. Data are mean (averaged across replicates) vst-normalized gene abundance.

**Figure 6.**
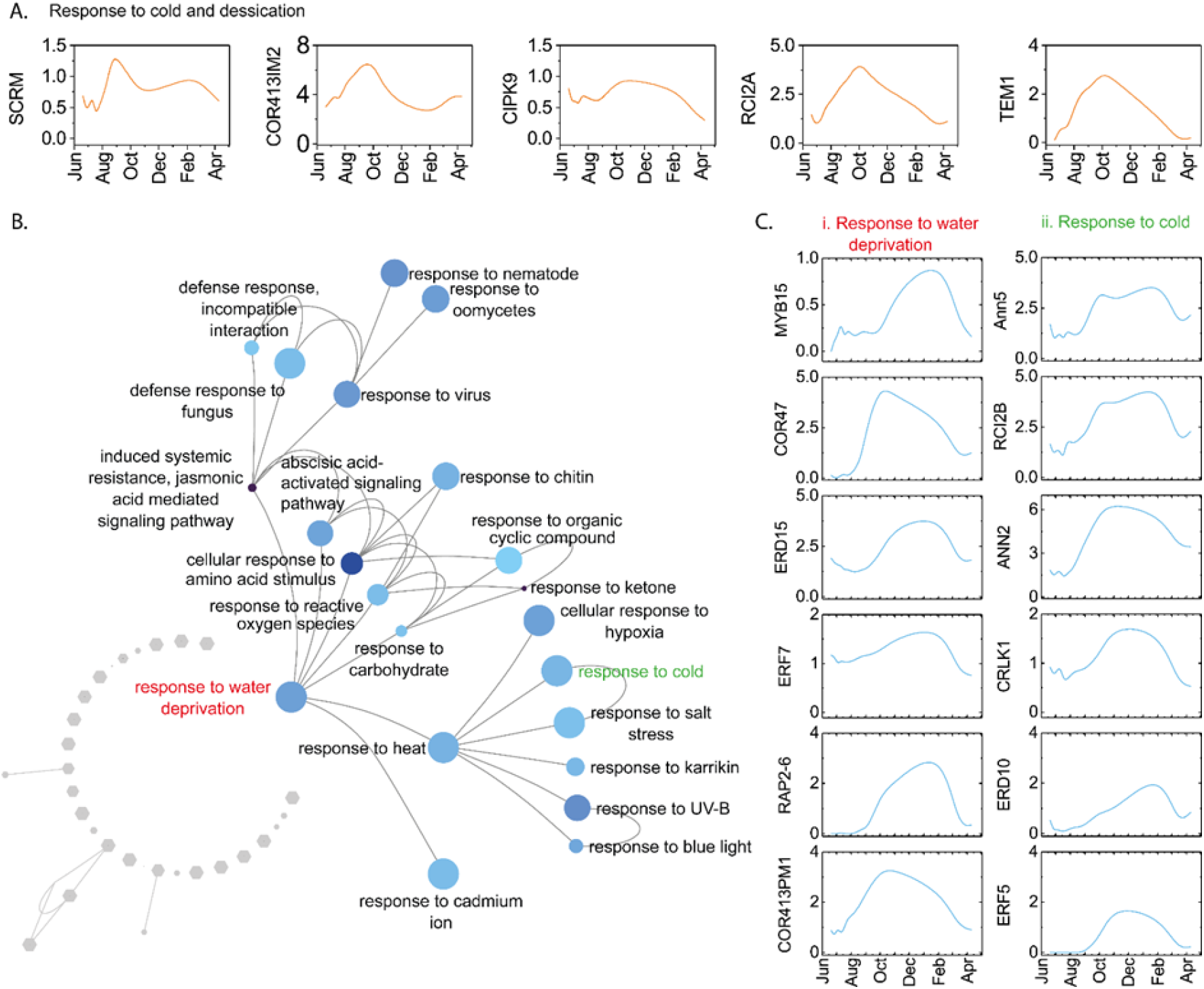
Gene Ontology (GO) term enrichment analysis of the genes in the blue cluster (IV) upregulated in winter and the temporal patterns of selected genes involved in desiccation and cold responses. ◻ A. The expression patterns of cold and desiccation related genes encoding SCRM (ICE1), COR412IM2, CIPK9, RCI2A and TEM1 from the orange cluster (I) upregulated in autumn. B. Enriched GO terms for biological processes in cluster IV upregulated in winter, with a major focus on nodes connected (Direct or indirect) to response to water deprivation. Size of the node is proportional to the (−)log_10_(P) value; larger node size represents higher significance and darker colour indicates higher fold enrichment of the GO term; C. The expression of genes encoding MYB15, COR47, ERD15, ERF7, RAP26 and COR413PM1 associated with response to water deprivation (C-i), and ANN5, RCI2B, ANN2, CRLK1, ERD10 and ERF5 associated with response to cold (C-ii). Data are mean (averaged across replicates) vst-normalized gene abundance.

Response to water deprivation was the most central node in the GO network of the gene cluster with enhanced expression in winter (Fig. 6B). Genes encoding cold and drought responsive proteins such as ethylene response factors (ERFs) and early response to dehydration, ERD15 and ERD10, showed slightly different dynamics over the winter (Fig. 6C i, ii). In addition, we found enhanced expression of TEM1 (TEMPRANILLO 1) related to ethylene signalling and to photoperiodic regulation already in the autumn (Fig. 6A). Several genes responsive to cold were also upregulated already in the autumn (cluster II, Fig. 2 and 4) such as ICE1 (Inducer of CBF expression 1 or SCRM), COR413IM2 that encodes thylakoid a membrane protein supposed to provide freezing tolerance, and calcineurin B-like (CBL)-binding protein kinases CIPK9, CIPK12 and CIPK26 (Fig. 6a). CBL proteins act as calcium sensors and they can be the link to activated calcium signalling in winter (Fig. 4). These results are consistent with needles responding to the decreasing photoperiod and starting to initiate winter acclimation already in September and October.

During the spring, the combination of low temperatures and high irradiation leads to high photooxidative stress in needles (see e.g. Bag et al 2020a). Not surprisingly, genes homologous to UV/high light responsive genes such as PHR1 (Phosphate Starvation Response 1), ELIP1 (Early light induced proteins), RUP3 (Repressor of UV-B Photomorphogenesis 3) and transcription factor MYB4 were upregulated (Table S4). In addition, the expression of nitrate transporters was elevated in spring (Fig. 4, Cluster I) perhaps reflecting reactivation of nutrient acquisition and growth. Spring and summer showed common upregulated processes such as cell wall modification (BXLs beta-xylosidase like proteins that are involved in pectic arabinan modification, cellulose synthases and callose synthases) and long-chain fatty-acyl-CoA metabolic process (FARs), presumably related to cuticle formation (Fig. 4, Cluster V, Table S4,S5).

GO enrichment analysis of genes, by comparing each consecutive season, was also performed to visualize the molecular function and cellular component (Figure S4-S10). GO enrichment analysis of season specific DE genes depicting the biological processes, molecular function and cellular component are shown in Data S1.

### Role of transcription factors in seasonal adaptation of cellular functions

This dataset provides us with an overview of the seasonal adaptation that occurs in Norway spruce needles to adapt to varying conditions and challenges. To further elucidate how transcription factors (TFs) were involved in each seasonal adaptation we performed DESeq analysis by comparing one season to each of the others. We identified several transcription factor families affected by specific seasons (Fig. 7, Fig S3, S4, Table S6) and in the following we will show examples from some of the most affected TF families.

**Figure 7.**
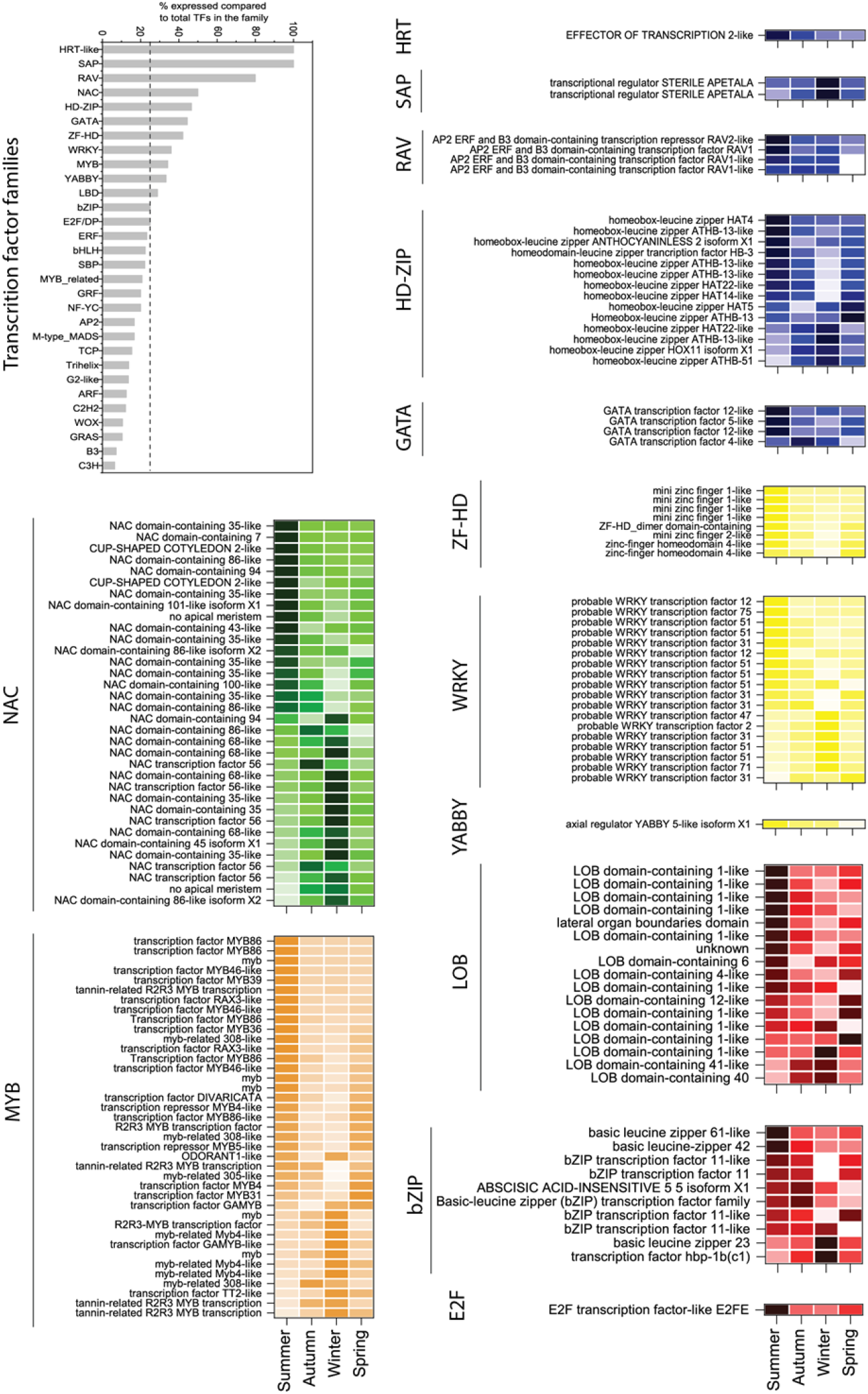
Season-specific expression of transcription factor families. ◻ Bar plot showing the percentage of expressed TFs by TF family, and heatmaps illustrating expression patterns of transcription factors in families in which more than 25% of the family members were differentially expressed in a season-specific manner (comparing each season against all other seasons in DESeq2, log_2_ fold change >2.0, FDR-adjusted *P*-value <0.01). Darker shading indicates greater fold-change of differential expression (DE). See complete list of DE gene lists Suppl. Table 6.

The NAC transcription factor family consists of NAM, ATAF and CUC transcription factors and out of 68 genes assigned to this family in spruce more than half (35) were differentially expressed during the year (Fig. 7). NAC family transcription factors are known to regulate a multitude of developmental processes such as senescence, cell division and wood formation as well as plant immunity and stress responses. Many of the 15 NAC TFs that were upregulated in summer were similar to those related to xylem formation and drought and/or salt stress resistance (Fig. 7). Six senescence associated or stress responsive NAC TFs (homologous to AtATAF1/NAC002, NAC025, NAC028, NAC086) were upregulated in autumn and eleven were induced in winter.

Out of 114 TFs belonging to the MYB TF family, 42 were differentially expressed during the year. In summer, highly expressed MYB TFs were similar to those related to cell wall biosynthesis (xylem formation, lignification), flavonoid biosynthesis and phenylpropanoid biosynthesis. In in winter, some upregulated MYBs were instead similar to those linked to cold and freezing tolerance (MYB15, Fig. 6c-i), UV-B response, growth arrest and anthocyanin biosynthesis. Almost 30% of WRKY family TFs (18 out of 50) were found to be differentially expressed. WRKY TFs related mostly to phosphorus related stress and phosphorus signalling were upregulated in summer, whereas in winter some WRKY TFs similar to those related to regulating plant innate immunity and salicylic acid (SA) and jasmonic acid (JA) crosstalk were upregulated.

The BHLH (Basic helix loop helix) family is the third largest TF family in spruce and the expression of 24 genes out of total 107 was significantly affected by season (Figure S3). In summer, among upregulated BHLH TFs were some similar to those related to abiotic stresses such as drought, salinity, and iron deficiency. In winter, most upregulated TFs were linked to phytochrome signalling.

Almost 50% of HD-ZIP TFs were affected by season. During the summer months different growth, sucrose signalling, accumulation of anthocyanin and root development related TFs were upregulated, whereas in winter growth inhibition and cold response related TFs were upregulated. Several GATA and ZF-HD TFs were also differentially expressed: in summer these were mostly related to light responsive functions and hormone signalling, respectively. Apart from the TF families mentioned above, transcription factors in, for example, LOB, bZIP, HRT, SAP, RAV, E2F and YABBY families were also differentially expressed (SI 4).

## Discussion

The ability of northern boreal conifers to adapt and survive harsh long winters without shedding their needles indeed is an extreme example of survival strategies in the plant kingdom. In this study, we performed deep RNA sequencing to provide a high-quality dataset useful to decipher regulation of seasonal acclimation of the cellular biochemical and metabolic activity triggered by changes in nuclear transcription. We show that needles collected over the annual cycle could be broadly assigned into five groups, representing early summer, summer, autumn, winter, and spring based on their transcriptome profiles. Overall, developmental processes were highly active in early summer and the acclimation processes that were expressed later in summer were in general inactivated in winter and vice versa.

As conifer genomes are sequenced at an impressive speed and tools such as genetic transformation of conifers are slowly becoming more accessible, it is likely that conifer biology will become increasingly interesting for scientists, given the enormous ecological and economic importance of these species. To date few extensive conifer RNA-Seq data sets are freely available and easily accessible, and we believe that our dataset, comprising of several samples from the same tissue and covering the whole year could represent a unique asset. To demonstrate the usefulness of the dataset we show that meaningful information on both the timing of different developmental processes in young needles, and acclimation in mature needles, can be extracted from the data. Moreover, we have shown that the analysis and interpretation of data not only can be conducted in multiple ways (differential expression performed with one season comparing to the other seasons, see Data S1 and S2), but also that different analyses for specified time periods reveal different phases of plant development and acclimation in more detail (Fig. 2, Figure S2, Table S2,S3). Samples collected, for example, in early summer overpower the variation of weather dependent transcriptome changes during the other seasons, due to the overall higher variation in gene expression levels in the former samples, combined with an extensive sampling during this period, motivated by a wish to capture many developmental processes taking place in a short time period. However, the analysis of early summer samples separately shows how the development of new needles is matched to transcriptome changes, whereas the analysis of transcriptome changes across summer to spring provides more information on the seasonal acclimation. We have avoided drawing conclusions from the expression profiles of individual genes but provide evidence, by analysing several families of transcription factors, that the dataset we provide provides high-quality information also at the level of the single gene, and therefore could regarded to be a transcriptome atlas of Norway spruce needles, made available for scientific community.

Although we believe that the main value of this contribution is to provide a resource to the other researchers, we also consider it useful to summarize the knowledge we have added to answer the question: what do Norway spruce needles actually do over the year? In Figure 8 we propose a hypothetical, even speculative, model representing the regulation of cellular activities in needles throughout the year. The five seasons have been given different colours and the outer circular bars indicate the main active cellular processes during the periods. We have divided them into five main modules of cellular activity, 1) Primary growth, 2) Secondary growth, 3) Response to biotic stimuli, 4) Winter stress related processes and 5) Response to abiotic stimuli. Furthermore, within the five main modules we show several subgroups and the specific active timing of these subgroups are also shown. In every module, subgroups of activity are shown with the colour of the main module, where darker shades indicate the times when that activity is the highest.

**Figure 8.**
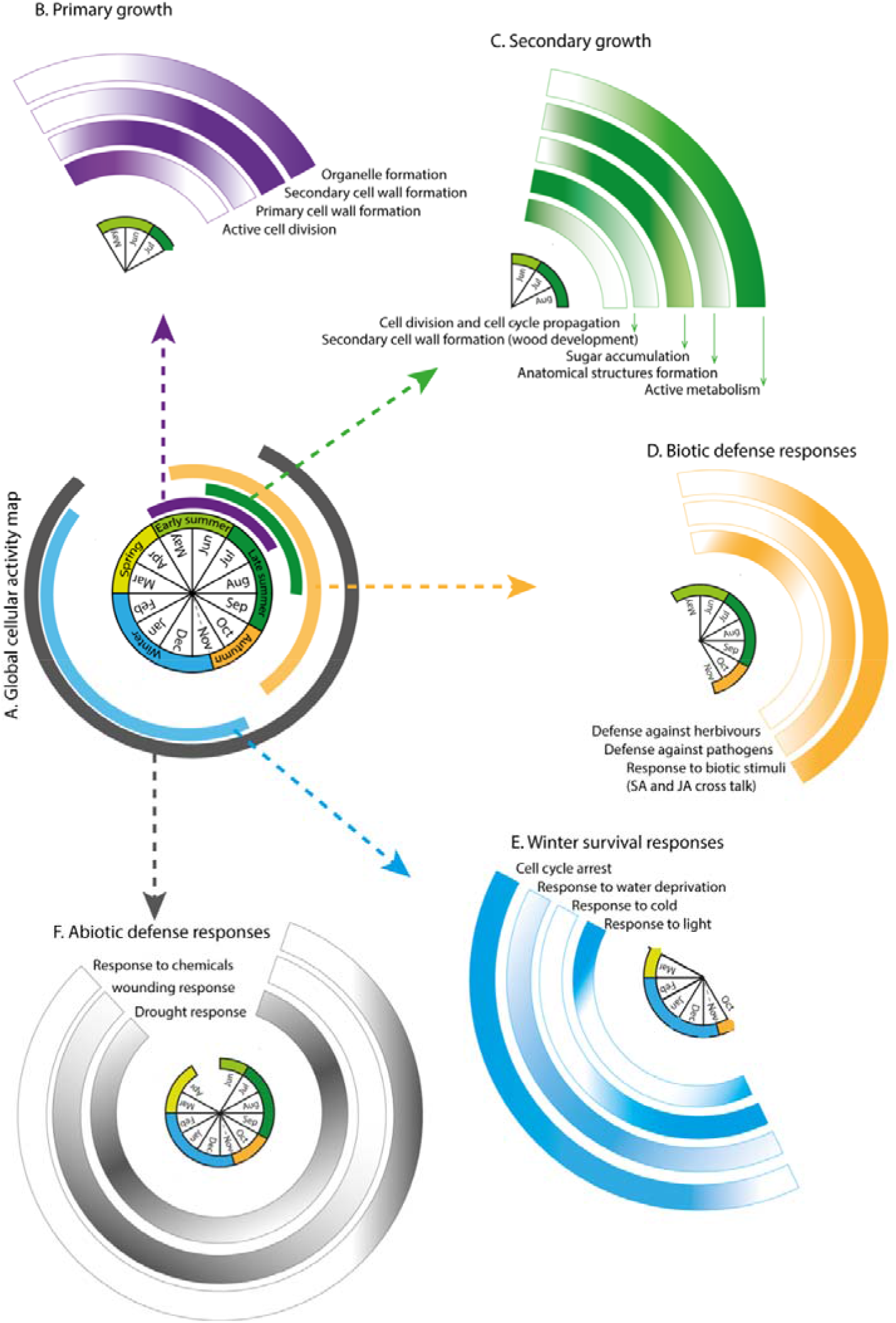
Hypothetical model based on RNA sequencing analysis representing seasonal acclimation of major cellular activities in Norway spruce throughout the annual cycle. The arcs represent the activity of genes (related to the functions) during the time period, where darker colour indicates higher and transparent means lowest activity. This map is a mere hypothetic representation of the overall cellular activity in spruce needles during one growth year and does not correspond to any empirical values.

In respect to primary growth and development (Fig. 8B), four major processes were largely affected by the seasonal cycle: cell division, primary cell wall formation, secondary cell wall formation, and organelle formation. Cell division is highly active at the beginning of early summer (May), declining towards June. Cell cycle, cell growth, DNA binding and replication were enriched GO terms in early summer and this period is accompanied with relatively high temperatures and long days suitable for growth, given a high photosynthetic activity in older leaves, and developing activity in current-year needles (Reich et al. 1995; Sveshnikov et al. 2006; Ivanov et al. 2002) as reported previously (Rossi et al. 2006). Genes involved in carbohydrate metabolism are highly expressed during summer and photosynthesis drives primary and secondary cell wall formation immediately in young needles, and growth in the rest of the tree. Cell wall deposition in needles and the formation of organelles take place in late June (Fig. 8B) concomitant with the high expression of anthocyanin and other photoprotective pigments, accompanied by expression of related transcription factors (Such as UV-B, HD-ZIP family). Secondary cell wall carbohydrate metabolism genes and related transcription factors (MYB and NAC family) are highly expressed in late summer (Fig. 8C), and wood formation and xylem lignification(Hartmann et al. 2017) are positively affected by cell wall related carbohydrate metabolism, which is elevated during this period (Jokipii-Lukkari et al. 2018). Other active processes are lipid and secondary metabolite formation followed by cell encapsulation, cuticle development, completion of internal organ development and the maturation of the needle.

Genes involved in biotic stress responses - signal transduction, cellular homeostasis, and cell communication (Fig. 4) - are mainly expressed from May to September (Fig. 8D). Signal transduction includes activity of kinases (Volkov 2012) and high enrichment of kinase activity (broadly as catalytic activity) is seen in the molecular function enrichment (Figure S4) and biological process analysis (Fig. 4, 8D). Cellular homeostasis GO terms follow a similar seasonal pattern to GO terms for high biotic stress responses and cell to cell communication. With respect to biotic stress, two main processes predominate: herbivory response in early summer and salicylic acid and jasmonic acid cross-talk from May until September.

We noted that the cell cycle arrest specific GO term was enriched in the winter module (Fig. 8E) when resources are allocated to survival, not growth. Potentially the resources must be recycled in winter (via salvage pathways) to fuel metabolic activities. In addition, plastid RNA modification was downregulated during winter accompanied by overall repression of transcription. Response to cold related activities such as upregulation of CBF factors and regulators were observed already in autumn (ICE1/SCRM) and other TFs (MYB and BHLH TFs) and genes encoding cold responsive (COR) proteins potentially accounting for freezing tolerance were highly expressed in winter (Miura and Furumoto 2013). We also found several NAC family transcription factors similar to those responding to water logging and hypoxia (Lee et al. 2014; Mancuso and Shabala 2010) highly expressed in winter (Fig. 7). We and others have shown that the photosystems are severely quenched in winter (Ottander, Campbell, and Öquist 1995; Ivanov et al. 2002; Öquist and Huner 2003; Bag et al. 2020b) and carbohydrate metabolism is supposedly minimal (Fig. 4, 6). Light and UV responses were upregulated during autumn and later in spring, following the photoperiod, temperature and solar radiation patterns during this time of the year (Fig. 1A). Finally, and somehow overlapping with winter survival responses, response to abiotic stimuli seems to be highly active throughout the year (Fig. 8F).

Taken together, our data and this model illustrate how Norway spruce needles shift between activity in the summer and survival responses in the winter as seen by the large number of differentially expressed genes during the transition periods, from autumn to winter and from winter to spring. As the transcriptome profiles seem to match well with what could be expected to be “required” in the needles, gene expression is important for the study of acclimation of spruce needles and our dataset is useful to examine it. Together with the visualization and analysis tools available in ConGenIE (tools such as GBrowse, BLAST and enrichment analysis tool, see Figure S4-S10, Data S1 for representations obtained from ConGenIE enrichment tools), we believe that our dataset could be an important asset for researchers interested in conifer biology and/or seasonal adaption in general. The dataset can be used to complement controlled experiments or be integrated in meta-analyses to decipher the molecular mechanisms underlying seasonal acclimation strategies such as early cold acclimation in autumn, freezing tolerance in winter and growth recovery in spring that all are fundamental for the survival of conifers in boreal regions. Important transcription factors can be identified and studied with, for example, genome editing approaches.

To conclude, in this contribution we present a comprehensive dataset on gene expression in Norway spruce needles over the year, and the database with all data and many analysis tools is available for the scientific community in ConGenIE (Sundell et al. 2015).

## Experimental procedures

### Plant material and study design

During the 2011-2012 growing season Norway spruce needles were harvested 28 times from five clonal copies of individual trees (40+ years old, 8-10 meter tall mature trees) growing at the Skogforsk research station, Sävar, Sweden, at midday (10:00 - 12:00) for mRNA extraction and sequencing (see details of sampling dates in Table S1). Approximately 1-2 g of needles were collected from three separate south facing branches (lower, middle and top in reachable range, approximately 2 metres from the ground) of a single tree and in total five different trees were sampled.

Daily solar radiation (SR) and air temperature (T) are depicted in Fig. 1A. The weather data was obtained from the weather station at the Department of Applied Physics and Electronics, Umeå University (http://www8.tfe.umu.se/weatherold).

### RNA extraction and sequencing

Total RNA was extracted from 0.5 g tissue using the CTAB method (Wang and Stegemann 2010). Precipitated RNA was further purified using an RNeasy Mini Kit (Qiagen, Hilden, Germany) according to the manufacturer’s protocol. RNA concentration and purity were measured using a NanoDrop 2000 spectrophotometer (NanoDrop Technologies, Wilmington, DE, USA) and its integrity was analysed on an Agilent 2100 Bioanalyzer (Agilent Technologies, Waldbronn, Germany). RNA sequencing was performed in SciLife labs, Stockholm, Sweden.

### Pre-processing of RNA-Sequencing data

An overview of the analysis pipeline is presented in Figure S1. The data pre-processing was performed following the guidelines described in Delhomme et al. 2014(Delhomme et al. 2014). Briefly, the quality of the raw sequence data was first assessed (FastQC v0.10.1), and residual ribosomal RNA (rRNA) contamination was removed (SortMeRNA v2.1b)(Kopylova, Noé, and Touzet 2012). Data were then trimmed to remove adapter sequences (Trimmomatic v0.32(Bolger, Lohse, and Usadel 2014)), after which another quality control step was applied to ensure that no technical artefacts were introduced during the pre-processing steps. Read counts were quantified by Kallisto (v0.43.0)(Bray et al. 2016) by using *Picea abies* v1.0 cDNA library as a reference. An overview of the data processing steps and details of the RNA data processing method and parameters are available in the GitHub repository (https://doi.org/10.5281/zenodo.494989).

For the data quality assessment and visualization, the counts were normalized using the variance stabilizing transformation (VST) as implemented in the Bioconductor DESeq2 package (v1.16.1(Love, Huber, and Anders 2014)). The biological quality control - i.e. the similarity of the biological replicates - was assessed by Principal Component Analysis (PCA) and hierarchical clustering analysis (HCA) in R (version 4.0.0)(R-Core-Team 2015).

### Differential expression analysis

Genes that were differentially expressed during the year were determined using the DESeq2 package in R(Love, Huber, and Anders 2014). Genes with absolute log_2_ fold change (Lfc2) >2.0 and false discovery rate (FDR) adjusted *P*-value of <0.01 were considered differentially expressed. The main reason for choosing Lfc2 threshold was that the number of expressed genes (the library size) was lower in winter compared to other seasons and choosing a high cut-off reduces the effect of the smaller library size. The differential expression tests were performed two ways: to identify season-specific genes by comparing one season against all other seasons, and to identify genes that were differentially expressed between consecutive seasons. Details of the DESeq2 analysis are available in GitHub (https://github.com/UPSCb/spruce-seasonal-needles).

### Global genome patterns and hierarchical clustering analysis

Principal component analysis (PCA) was performed to study the variation in the global genome profile throughout the year using SIMCA-P+ (version 15, Umetrics, Sweden). Genes that were present in >80% of the samples were included in the analysis. The missing value threshold was increased from 50% (default) to 80% to reduce the interference of missing values on the PCA model fitting(Jackson 2005). The NIPALS algorithm was used to correct for missing values in the data in SIMCA-P+(Wold 1968). The data were log_10_-transformed and scaled with unit variance.

Hierarchical clustering analyses (HCA) were performed in three ways: with all data, with early summer data only, with differentially expressed (DE) genes between consecutive seasons, and separately with differentially expressed (DE) genes between early summer and other seasons, to identify clusters with distinct temporal patterns, using ggplot2 (package version 3.0.3, R version 4.0.0 (Wilkinson 2011)). HCA was based on the Pearson correlation coefficient (distance) with Ward’s linkage. Ward’s linkage method was applied to minimize the variance within clusters(Ward Jr 1963; Murtagh and Legendre 2014). The optimal number of gene clusters was determined with a cut-off of k=12 (global data), tree height of 20 (for early summer data) or tree height of 10 (for DE genes). Circular dendrograms were produced using the dendextend R package (version 1.13.4, (Galili 2015)) in R.

### Analysis of transcription factors

Transcription factors (TFs) were identified based on the list obtained from the Plant Transcription Factor Database (PlantTFDB, http://planttfdb.cbi.pku.edu.cn/)(Jin et al. 2014; 2015; Tian et al. 2020). Heatmaps displaying the expression patterns of the most enriched TF family members affected by the seasons were visualized with Origin Lab (Origin Pro 2017 version 9.4.2.380).

### Database and analysis tools

Gene Ontology (GO) term enrichment analyses for biological processes (SLIM GO), molecular function and cellular components were performed for gene clusters with Panther (http://pantherdb.org/(Mi et al. 2019)) using the best Blast2GO(Götz et al. 2008) hits for *Arabidopsis thaliana* gene IDs obtained from ConGenIE (https://congenie.org/). Only the enriched parent terms in each cluster were included in the seasonal GO term network that was produced with REVIGO (http://revigo.irb.hr/)(Supek et al. 2011) and visualized with Cytoscape (version 3.8.0). For REVIGO analysis, a medium (0.7) similarity cut-off was used and *Arabidopsis thaliana* was chosen as a model organism. Resnik’s similarity method(Resnik 1999) was chosen for similarity measures between GO terms. Log_10_ *P*-values were provided alongside GO term ids as a measure of significance of the particular GO term.

## Accession number/Data availability

All data presented in the main manuscript or in supporting information are provided in either along with the figures S1-S10, tables S1-S6, data S1-S2 or detailed codes and raw values are given in github (https://github.com/UPSCb/spruce-seasonal-needles). The raw data is available from the European Nucleotide Archive (ENA) under accession PRJEB26453.

## Acknowledments

Members of the UPSC conifer genomics team – in particular Chanaka Mannapperuma, Thomas Hiltonen, Susanne Larsson and Simon Birve – are acknowledged for assistance in the preparation of samples and processing of data and Nathaniel R Street for support and comments on the manuscript. The computations were enabled by resources provided by the Swedish National Infrastructure for Computing (SNIC) at UPPMAX, partially funded by the Swedish Research Council through grant agreement no. 2018-05973 and SciLife lab provided both resources and comprtence. This work was supported by SE2B Horizon 2020 under grant agreement no. 675006 (SE2B), the Swedish Research Council VR, FORMAS, the Swedish Research Council for Environment, Agricultural Sciences and Spatial Planning, the Knut and Alice Wallenberg Foundation, the Swedish Governmental Agency for Innovation Systems (VINNOVA), the Kempe foundation, and the Trees for the Future (T4F) project.

## Author Contributions

SJ and PB conceived the idea; ND and TR processed the transcriptome data, PB, ND and JL designed and performed the gene expression analysis, PB, ND, JL, KMR and SJ integrated the results and PB, JL, KMR and SJ wrote the paper.

## Notes

**Conflict of interest:** Authors declare no conflict of interests.

### Competing Interest Statement

The authors have declared no competing interest.

